# Predictive attenuation of touch and tactile gating are distinct perceptual phenomena

**DOI:** 10.1101/2020.11.13.381202

**Authors:** Konstantina Kilteni, H. Henrik Ehrsson

**Affiliations:** Department of Neuroscience, Karolinska Institutet, Solnavägen 9, 17165 Stockholm, Sweden

## Abstract

In recent decades, research on somatosensory perception has led to two important observations. First, self-generated touches that are predicted by voluntary movements become attenuated compared to externally generated touches of the same intensity (attenuation). Second, externally generated touches feel weaker and are more difficult to detect during movement than at rest (gating). Currently, researchers often consider gating and attenuation the same suppression process; however, this assumption is unwarranted because, despite more than forty years of research, no study has combined them in a single paradigm. We quantified how people perceive self-generated and externally generated touches during movement and rest. We show that whereas voluntary movement gates the precision of both self-generated and externally generated touch, the amplitude of self-generated touch is robustly attenuated compared to externally generated touch. Furthermore, attenuation and gating do not interact and are not correlated, and we conclude that they represent distinct perceptual phenomena.

## Introduction

Let us imagine you are at your doctor for a medical examination. Upon her request, you apply pressure with your index finger to your leg to indicate exactly where you feel the pain. The pressure you feel on your leg and the tip of your finger is feedback from your voluntary finger movement, and it is called *somatosensory reafference*. Imagine that the doctor now applies pressure with her index finger to the same spot on your leg to reproduce and confirm your sensations. This pressure is generated by the doctor and not by you and it is called *somatosensory exafference*. Now, imagine that the doctor asks you to first keep your leg relaxed and then flex and extend it while you or her continuously apply pressure to your leg. You therefore experience your reafferent or her exafferent touches on your leg while it is moving or resting. Distinguishing between these four conditions is fundamental for your sensorimotor control; your nervous system must know both the source of the touch and the state of your limb to appropriately use the sensory feedback. A cutaneous mechanoreceptor in your peripheral nervous system, however, is unable to distinguish whether a touch is reafferent or exafferent, and thus, this distinction must be made centrally, where tactile signals from the skin, sensory information from muscles and joints, and information from motor commands are available. How, then, does the central nervous system classify somatosensory signals during movement?

Several experimental studies in humans have shown that the brain *attenuates* somatosensory reafference (i.e., all somatosensory inputs generated by one’s own movement, including inputs originating directly from the moving body part and inputs from other passive body parts being touched) compared to exafference. In behavioral research, this process refers to participants perceiving self-generated strokes, forces, or taps as weaker than external equivalents of the same intensity (*1, 2, 11, 3*–*10*). This somatosensory attenuation is related to reduced activity in the secondary somatosensory cortex (*12*–*14*) and the cerebellum (*12, 14, 15*) and increased connectivity between the two areas (*14, 16*) during self-generated touches compared to externally generated touches. Somatosensory attenuation was observed in 98% or 315 of 322 people across a wide age range (*17*), and it is considered one of the reasons why we cannot tickle ourselves (*18*–*20*).

Sensory attenuation is not exclusive to humans; similar strategies are used by other species across the animal kingdom (for reviews, see (*21*–*25*)). For example, during self-chirping, the cricket’s central auditory processing is inhibited (both presynaptically and postsynaptically) in phase with the insect’s chirps to prevent desensitization of its auditory system while maintaining sensitivity to external sounds (*26, 27*). In mice, auditory cortical responses to self-generated sounds are attenuated, and this attenuation is present only for the tone frequencies the animal has associated with its locomotion and absent when the same sounds are externally produced (*28, 29*). A weakly electric fish (*30*) is able to respond exclusively to externally generated electrical discharges by attenuating its predicted electrosensory reafference (*22, 31*). In primates, activity in the vestibular nucleus in response to vestibular reafference is attenuated during active head movements compared to passive head movements, allowing the animal to maintain its head and body posture and activate vestibular-related reflexes when appropriate (*21, 22, 32*–*34*).

At the same time, another branch of experimental research has shown that somatosensory sensitivity in response to externally generated stimuli is *gated* during and before a voluntary movement. In human psychophysical research, this phenomenon of *movement-related tactile gating* or *tactile suppression* manifests as an increase in the detection threshold (*35*–*40*), a decrease in the detection rate (*35*–*37, 41*–*45*), a decrease in the detection precision (*38, 40, 46, 47*), and a decrease in the subjective intensity of externally generated stimuli (*37, 42, 48*) when the stimulated body part moves compared to when it is at rest. Several electrophysiological studies have shown that this gating reflects inhibition of somatosensory evoked potentials during active movement compared to rest at subcortical and cortical sites along the somatosensory pathway (*48*–*53*). Similar to somatosensory attenuation, tactile gating is a biologically preserved mechanism that is observed across different species (*54*). For example, responses recorded in the cat medial lemniscus evoked by nerve stimulation are suppressed prior to and during limb movements (*55*). Similarly, the transmission of cutaneous afferent signals to the primary somatosensory cortex is suppressed in rats during movement compared to rest (*56*). In monkeys, the gating of cutaneous afferent input during active movement has been observed in both the primary somatosensory cortex (*57*–*59*) and the spinal cord (*59, 60*).

Somatosensory attenuation and tactile gating share two important conceptual similarities. First, they both refer to modulation, either in terms of magnitude or precision, of the perception of cutaneous stimuli during movement. Second, they have been assigned the same functional role (*43*): to reduce the flow of afferent information that can be predicted from the motor command and enable the detection of an external input that may be biologically important, such as touches caused by predators (*18, 25, 61*), or input that is task-relevant for the upcoming or ongoing movement (*50, 52, 62*).

Importantly, however, the two phenomena present one striking difference. Somatosensory attenuation relates to somatosensory reafference, that is, touches caused *by* our voluntary movement. In contrast, gating relates to somatosensory exafference, that is, external touches occurring *during* our voluntary movement. Nevertheless, somatosensory research often treats the two phenomena as a single generalized suppression strategy of the brain. For example, reviews and theoretical papers use the terms attenuation and gating (*63, 64*) or their literature (*65*) interchangeably. This intermix is also evident in experimental studies and more specifically in the design, measures, and interpretation of the findings. For example, some experiments have tried to relate neural responses triggered by externally generated touches (i.e., electrical stimulation) to the responses associated with self-generated touches (*66, 67*), or they applied externally generated touches during a self-generated movement and interpreted them as self-generated touches (*68*). Other studies have applied externally generated touches (*38, 39*) but used theories for the perception of self-generated touches to explain their findings.

If the two phenomena are indeed different, this false equivalence is detrimental for our understanding of human sensorimotor control. First, it prevents advances in our understanding at the computational level because researchers try to explain gating (e.g., (*38*)) and attenuation (e.g., (*69*)) using the computational processes proposed for attenuation and gating, respectively. Similarly, at the neurobiological level, researchers intermix neural correlates of gating and attenuation (e.g., (*66, 67*)) because they assume that they measure the same single phenomenon. Second, this confusion becomes particularly disadvantageous in clinical studies using gating and attenuation when interpreting findings of sensorimotor deficits in patients with schizophrenia (*70*–*72*), functional movement disorders (*73*) or Parkinson’s disease (*50, 74, 75*).

Are these phenomena the same, or do they represent two fundamentally distinct processes? Does the brain treat all sensory stimuli similarly during movement, regardless of whether they are reafferent or exafferent? To the best of our knowledge, no previous study has attempted to simultaneously test attenuation and gating with the same stimulus and psychophysics task. Here, in a single experimental design, we investigated the perception of touches applied on the left hand while manipulating whether the left arm was in movement or resting (left limb state). We additionally manipulated whether the touches were reafferent, generated by the right hand, or exafferent, generated by an external source (origin of touch). We reasoned that if the two phenomena are the same, they should influence somatosensory perception in similar manners. Our results do not confirm this hypothesis: voluntary movement reduces the somatosensory precision of both sensory reafference and exafference (gating), while the perceived amplitude of sensory reafference is robustly attenuated compared to that of exafference (attenuation). Notably, the two phenomena do not correlate with each other or interact when present simultaneously. Thus, collectively, our results show that gating and attenuation are two separate processes that can be experimentally dissociated.

## Results

Participants rested their left hands palm up with their index fingers on a molded support, and their right hands were placed palm down on top of a set of sponges (**Fig. 1a-d**). In all conditions, they performed a force-discrimination task (*3, 4, 10, 71*): in each trial, a motor delivered two taps (the *test* tap and the *comparison* tap) on the pulp of their left index finger, and they were asked to verbally indicate which tap felt stronger (**Fig. 1e-h**). While the *test* tap had a fixed intensity (2 N), the intensity of the *comparison* tap randomly changed in every trial (1, 1.5, 1.75, 2, 2.25, 2.5, or 3 N). An auditory ‘go’ signal indicated the trial onset and the onset of the response period.

**Fig. 1.**
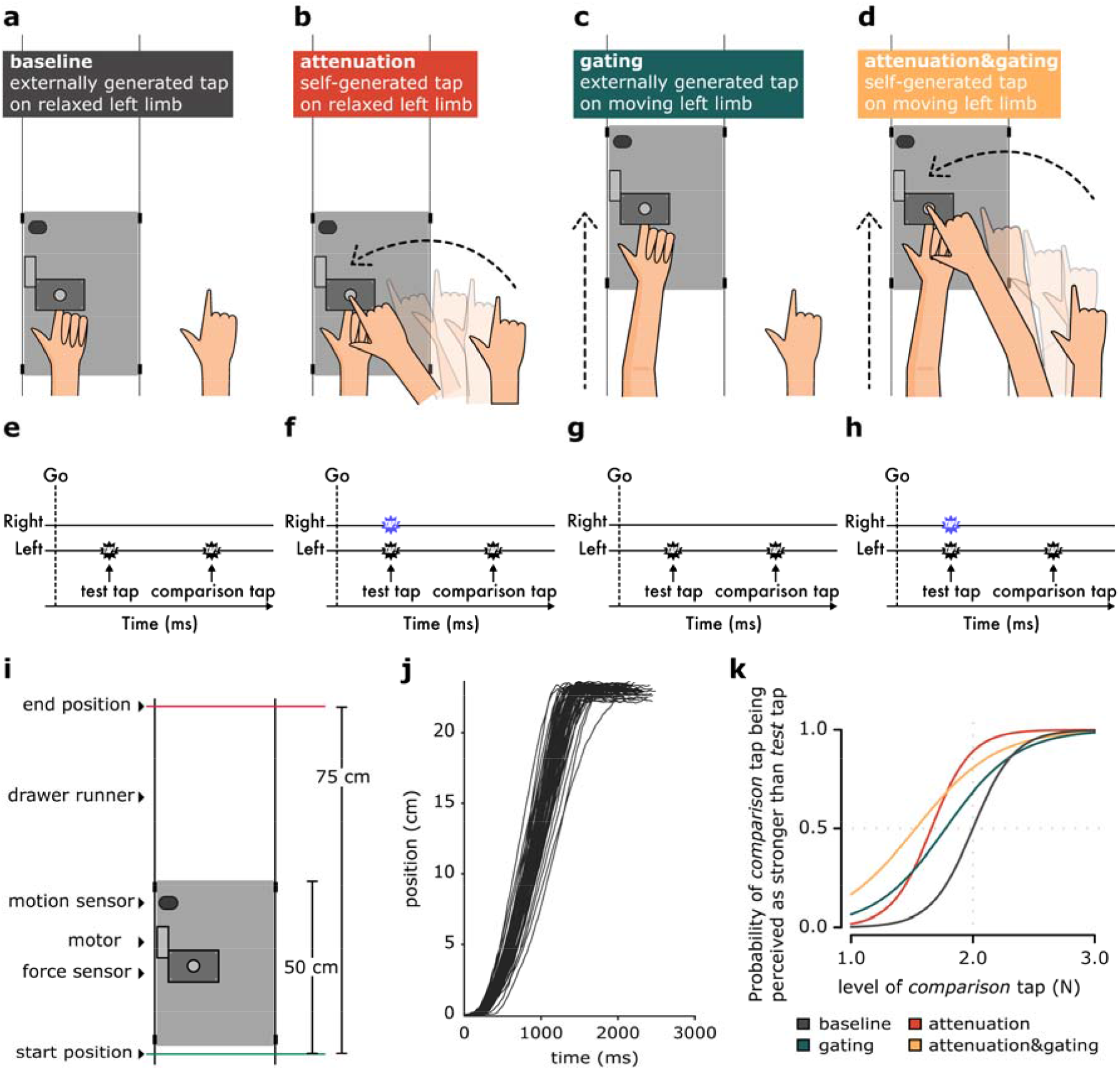
Experimental conditions. Two factors were manipulated in the experiment, resulting in four experimental conditions: whether the left arm was in rest (**a, b**) or in movement while receiving the *test* tap (**c, d**) (state of the left limb) and whether the *test* tap was externally triggered by the motor (**a, c**) or self-triggered by the participants’ right hand (**b, d**) (origin of touch). In all four conditions **(e-h)**, the participants received the two taps (*test* and *comparison* tap) on the pulp of their left index fingers from the motor (black stars), and they were required to verbally indicate which was stronger: the first or the second tap. In the *attenuation* and *attenuation&gating* conditions, the participants self-triggered the test tap on their left index finger by moving their right arm to tap a sensor with their right index finger (blue stars). **(i)** In the *gating* and *attenuation&gating* conditions (**c** and **d**), the participants extended their left arm from a starting position to an ending position, sliding the experimental setup along two drawer runners. During this movement, participants experienced the *test* tap. A motion sensor recorded the position of the platform in time. **(f)** Example position traces recorded by the motion sensor for the movements of one participant during the *attenuation&gating* condition. **(g)** Fitted logistic models of the responses of one example participant to the four experimental conditions.

In all conditions, our experimental manipulation exclusively concerned the *test tap*; the *comparison* tap was always externally triggered and delivered on the relaxed left arm, serving, therefore, as a reference stimulus. In a factorial design, we controlled for whether the left arm moved **(Fig. 1c, d)** or remained relaxed **(Fig. 1a, b)** during the *test* tap and whether the *test* tap was produced by the right hand (self-generated) **(Fig. 1b, d)** or not (externally generated) **(Fig. 1a, c)**. This design resulted in four experimental conditions, the order of which was fully counterbalanced (**Fig. 1a-d**).

In the *baseline* condition (**Fig. 1a**), participants did not move their limbs but passively received the *test* and the *comparison* taps on the left index finger. This control condition was used to assess the participants’ somatosensory perception in the absence of any movement (*2, 4, 10*). In the *attenuation* condition (**Fig. 1b**), participants actively tapped a force sensor placed on top of their left index finger with their right index finger. The tap of their right index finger on the force sensor triggered the *test* tap on their left index finger. This classic condition was used to assess the perception of a self-generated tap on a passive limb (*2, 4, 10*). In the *gating* condition (**Fig. 1c**), participants were asked to continuously move their left arm forward, sliding the experimental setup with the motor and the force sensors between a start and an end position (distance 25 cm) at a comfortable velocity of approximately 20 cm/sec (**Fig. 1i-j, Supplementary Fig. S1, S2**). During this movement, participants received the *test* tap on their left index finger. This condition represents gating because it assesses the perception of an externally generated tap on a moving limb(*41, 42, 44, 76*). Finally, in the *attenuation&gating* condition (**Fig. 1d**), the participants performed the same movement with their left arm, but they were additionally asked to actively tap with their right index finger on the force sensor that triggered the *test* tap on their left index finger during the movement. The force sensor was attached to the experimental setup and moved together with the left hand. This condition combines the gating and attenuation phenomena since it is used to assess the perception of a self-generated tap on a moving limb.

The participant’s responses to each condition were fitted with a generalized linear model (**Fig. 1k, Supplementary Fig. S3**). For all participants and all conditions, the fitted model was better than a null/restricted model with no predictors, according to McFadden’s R squared measure. Two parameters of interest were extracted: the point of subjective equality (PSE), which represents the intensity at which the test tap felt as strong as the comparison tap (*p* = 0.5), and the just noticeable difference (JND), which reflects the participants’ sensitivity (precision) in force discrimination. A lower PSE in an experimental condition indicates that the test tap felt weaker in that condition. A higher JND in an experimental condition indicates that the discrimination sensitivity was lower in that condition (i.e., a larger difference in the force intensities needed to be detected). PSE and JND correspond to different qualities of sensory judgments – accuracy and precision, respectively – and can be independent (*77*). We hypothesized that the two phenomena are different and, thus, they affect the PSE and JND differently. Specifically, we expected a decrease in the perceived magnitude (lower PSE) for conditions with sensory reafference (*attenuation* and *attenuation&gating* conditions), with no effects on the sensory precision of the participants (JND) when they received reafference on a still limb (*attenuation* condition). That is, a self-generated touch will feel weaker than an externally generated touch, but no effect on somatosensory precision will be observed. In contrast, we predicted a decrease in the somatosensory precision (higher JND) of both sensory reafference and exafference for conditions where the limb that receives the touches moves (*gating* and *attenuation&gating* conditions). That is, the precision with which a touch (either self-generated or externally generated) is perceived is lower on a moving limb because of the additional kinaesthetic, proprioceptive, tactile signals from the movement. A small, if any, effect on the PSE was expected for the *gating* condition. We hypothesized that the attenuation phenomenon mainly affects the PSE and not the JND, and the gating phenomenon mainly affects the JND and not the PSE. Our hypotheses were supported by the data (**Fig. 2a-g)**.

**Fig. 2.**
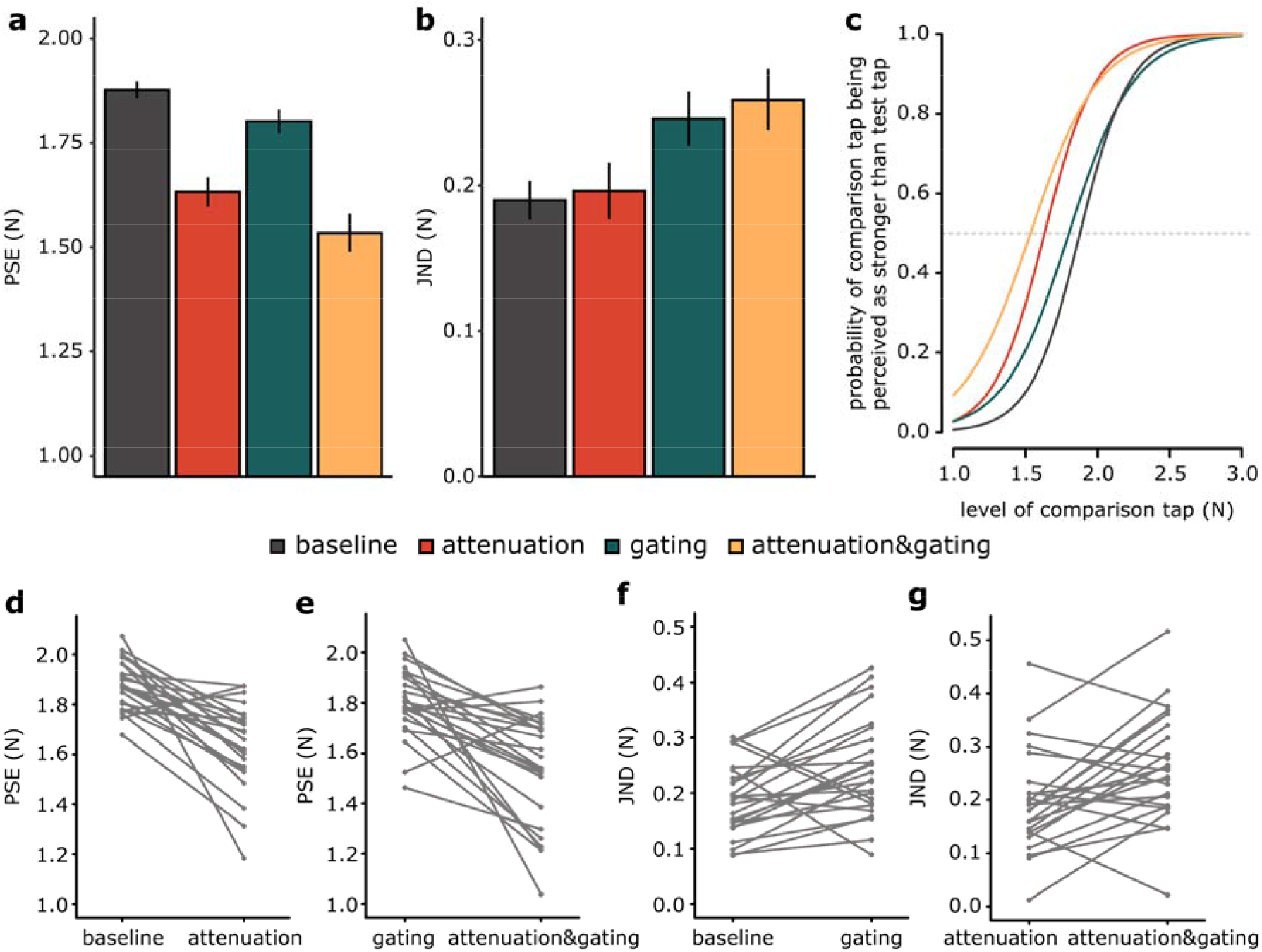
Experimental results. **(a, b)** Bar graphs showing the PSEs and JNDs (mean ± SEM) for each condition. A lower PSE value indicates a lower perceived magnitude, while a higher JND value indicates lower somatosensory sensory precision. Reafferent touches produced by the movement of the right arm (*attenuation* and *attenuation&gating* conditions) were associated with a significant decrease in the PSEs, while movement of the left arm that receives the touches (*gating* and *attenuation&gating*) results in a significant increase in the JNDs. A small decrease in PSE was also observed in the *gating* and *attenuation&gating* conditions compared to the *baseline* and *attenuation* conditions, respectively. **(c)** Group psychometric functions for each condition generated using the mean PSE and the mean JND across participants. The leftward shift of the curves for the *attenuation* and *attenuation&gating* conditions illustrates that somatosensory reafference is perceived as weaker than exafference. The flattening of the curves for the *gating* and *attenuation&gating* conditions illustrates the worsening somatosensory precision of both the reafference and exafference on a moving limb. **(d, e)** Line plots illustrating the decreases in PSEs when experiencing reafferent touches compared to exafferent touches when the left arm is still **(d)** and when the left arm moves **(e). (f, g)** Line plots illustrating the increases in JNDs when receiving touches on a moving limb compared to rest when touches are exafferent **(f)** and when touches are reafferent **(g)**.

We performed repeated-measures ANOVA on the PSEs with the origin of the touch (reafference *vs*. exafference) and the state of the left limb (movement *vs*. rest) as factors. This analysis revealed a significant main effect of the origin of the touch (*F*(1, 23) = 36.10, *p* < 0.001, *η*_*p*_^*2*^ = 0.611), a significant main effect of the left limb state (*F*(1, 23) = 13.91, *p* = 0.001, *η*_*p*_^*2*^ = 0.377) and a nonsignificant interaction (*F*(1, 23) = 0.26, *p* = 0.615, *η*_*p*_^*2*^ = 0.011) (**Fig. 2a)**. A Bayesian repeated-measures ANOVA further concluded against the interaction term by supporting the model without the interaction term compared to the full factorial model (*BF*_*M1*_ /*BF*_*M2*_ = 3.631). The *attenuation* condition produced a significant decrease in the PSE compared to the *baseline* condition (*n* = 24, *t*(23) = -5.908, *p* < 0.001, *Cohen’s d* = - 1.206, *CI*^*95*^ = [-0.332, -0.160], *BF*_*01*_ <0.0003) (**Fig. 2d)**. This replicates previous attenuation findings, indicating that a self-generated tap on a passive limb feels weaker than a tap of the same intensity but of an external origin (*2, 4*–*7, 10, 14, 17*). Similarly, the PSE in the *attenuation&gating* condition was significantly lower than that in the *gating* condition (*n* = 24, *t*(23) = -5.032, *p* < 0.001, *Cohen’s d* = -1.027, *CI*^*95*^ = [-0.377, -0.157], *BF*_*01*_ = 0.002) (**Fig. 2e)**, extending the previous conclusion to when the receiving limb is moving. Together, these two contrasts show that reafferent (self-generated) touches feel weaker than exafferent touches, both when the receiving hand is in movement and when it is at rest.

The *gating* and *attenuation&gating* conditions also resulted in a significant decrease in the PSE compared to that in the *baseline* condition (*n* = 24, *t*(23) = -2.409, *p* = 0.024, *Cohen’s d* = -0.492, *CI*^*95*^ = [-0.141, -0.011]) and the *attenuation* condition (*n* = 24, *V* = 55, *p* = 0.005, *rrb* = -0.633, *CI*^*95*^ = [-0.161, -0.022]), respectively. However, these decreases were quite modest (≌ 30% of the weakening produced by the *attenuation* condition) and supported only by anecdotal evidence from Bayesian statistics (*BF*_*01*_ = 0.433 and *BF*_*01*_ = 0.738, respectively). Together, these contrasts suggest that exafferent touches may feel slightly weaker on a moving limb than on a passive limb, consistent with previous findings regarding tactile gating (*37, 42, 48*). Nevertheless, compared with tactile reafference, the perceived magnitude of tactile exafference is not substantially decreased.

When testing for the effects of the conditions on the somatosensory precision of the participants (JND), a significant main effect of the state of the left limb was observed (*F*(1, 23) = 17.1, *p* < 0.001, *η*_*p*_^*2*^ = 0.426), but neither a significant main effect of the origin of touch (*F*(1, 23) = 0.52, *p* = 0.478, *η*_*p*_^*2*^ = 0.022) nor a significant interaction (*F*(1, 23) = 0.06, *p* = 0.809, *η*_*p*_^*2*^ = 0.003) was identified (**Fig. 2b)**. Similar to PSEs, the absence of an interaction for the JNDs was further supported by Bayesian repeated-measures ANOVA, which provided evidence against the interaction term (*BF*_*M1*_ /*BF*_*M2*_ = 3.522). The *attenuation* condition did not result in any change in the JND compared with that in the *baseline* condition (*n* = 24, *t*(23) = 0.331, *p* = 0.744, *Cohen’s d* = 0.068, *CI*^*95*^ = [-0.034, 0.047]), and this result was substantially confirmed by Bayesian analysis (*BF*_*01*_ = 4.432). Similarly, no significant differences in the JND were detected between the *gating* and the *attenuation&gating* conditions (*n* = 24, *t*(23) = 0.72, *p* = 0.481, *Cohen’s d* = 0.146, *CI*^*95*^ = [-0.024, 0.05]), which was again confirmed by the Bayesian analysis (*BF*_*01*_ = 3.691). Together, these two contrasts indicate that receiving sensory reafference *per se* is not accompanied by worsening of sensory precision on the receiving limb.

In contrast, moving the limb while receiving an external touch (*gating* condition) resulted in a significant increase in the JND compared to that in the *baseline* condition (*n* = 24, *t*(23) = 3.134, *p* = 0.005, *Cohen’s d* = 0.640, *CI*^*95*^ = [0.019, 0.093], *BF*_*01*_ = 0.108) (**Fig. 2f)**. This result was further confirmed by a significant increase in JND in the *attenuation&gating* condition compared with the *attenuation* condition (*n* = 24, *t*(23) = 2.984, *p* = 0.007, *Cohen’s d* = 0.609, *CI*^*95*^ = [0.019, 0.106], *BF*_*01*_ = 0.146) (**Fig. 2g)**. Together, these two differences indicate that voluntary movement *per se* decreases the precision with which reafferent and exafferent stimuli are perceived on the moving limb.

Together, our results indicate that predicting the sensory consequences of a voluntary movement produces a decrease in the perceived magnitude of sensory reafference (PSE) without a concomitant worsening of somatosensory precision (JND). In contrast, voluntary movement leads to a decrease in somatosensory precision (JND) for both sensory reafference and exafference. These effects are observed in the group psychometric fits (**Fig. 2c**). We tested whether we could better predict the participants’ performance in the *attenuation&gating* condition when using the PSE from the *attenuation* condition and the JND from the *gating* condition than the PSE from the *gating* condition and the JND from the *attenuation* condition, to further illustrate that somatosensory attenuation affects the amplitude (PSE) while tactile gating affects the precision (JND) and not *vice versa*. Indeed, the first model was significantly better: *n* = 24, *V* = 39, *p* < 0.001, *rrb* = -0.74, *CI*^*95*^ = [-68.980, -12.736], *BF*_*01*_ = 0.443 (**Supplementary Fig. S4, Fig. 3a-b**).

**Fig. 3.**
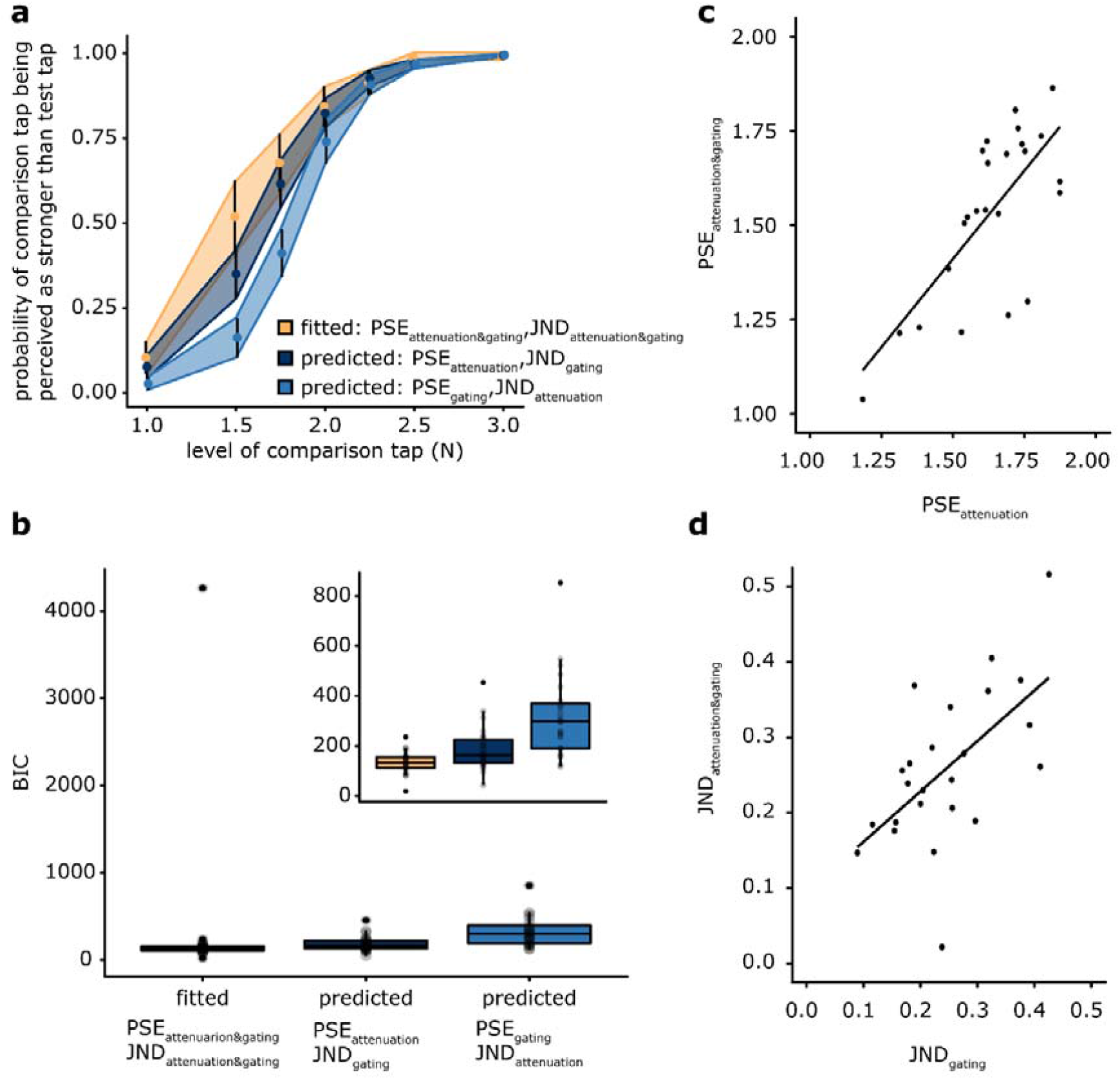
Model predictions and scatterplots for PSEs and JNDs. **(a)** Average responses of participants in the *attenuation&gating* condition (yellow) and average predicted responses using the parameters from the *attenuation* and *gating* conditions (blue). The responses depicted in dark blue indicate the PSE of the *attenuation* condition and the JND from the *gating* condition, while the responses depicted in light blue represent the PSE of the *gating* condition and the JND of the *attenuation* condition. The error bars and ribbons represent 95% confidence intervals. **(b)** For each participant, we estimated the Bayesian information criterion (BIC) of the fitted logistic model in the *attenuation&gating* condition and the two models with fixed parameters from the *attenuation* and *gating* conditions. The model using the PSE of the *attenuation* condition and the JND of the *gating* condition was a significantly better model than that using the PSE of the *gating* condition and the JND from the *attenuation* condition. The upper right panel presents the same data after excluding one participant corresponding to the outlier for illustration purposes. The exclusion of the outlier did not change the statistical results. **(c)** The participants’ PSEs in the *attenuation* condition were significantly correlated with those in the *attenuation&gating* condition. **(d)** The participants’ JNDs in the *gating* condition were significantly correlated with those in the *attenuation&gating* condition. No significant correlations between PSEs and JNDs were observed.

In the abovementioned ANOVAs, no significant interactions were observed between the two factors (the origin of touch and the state of the limb), neither for the PSEs nor for the JNDs, according to both frequentist and Bayesian analyses. Therefore, the effect of the left limb state was not influenced by the effect of the origin of touch, neither for the PSE nor for the JND. Instead, the two effects were summed when simultaneously present, and no superadditive effects were detected (interactions), indicating that each factor had its own independent effect on the responses. We performed a correlation analysis of the PSEs and JNDs to further test whether the effects produced by each phenomenon related to the others at all. No significant correlations were detected between any of the PSEs and any of the JNDs (all *p value*s > 0.225, *BF*_*01*_ = [1.971, 3.950]). The only significant correlation found for PSEs was between the PSE in the *attenuation* condition and the PSE in the *attenuation&gating* condition (*t*(22) = 4.89, *r* = 0.722, *p* < 0.001, *CI*^*95*^ = [0.449, 0.871], *BF*_*01*_ = 0.002) (**Fig. 3c**). That is, the weaker the participants perceived the magnitude of their self-generated touch during rest, the weaker the magnitude of their self-generated touch during movement felt. As the PSE values decreased significantly in these two conditions and these decreases correlated with each other, this result provides further support that their common experimental denominator, i.e., the reafferent nature of the touch, was responsible for the decrease in the PSE and the subsequent attenuation phenomenon. In contrast, the JND in the *gating* condition was significantly correlated only with the JND in the *attenuation&gating* condition (*t*(22) = 3.47, *r* = 0.595, *p* = 0.008, *CI*^*95*^ = [0.252, 0.805], *BF*_*01*_ = 0.047) (**Fig. 3d**). This specific correlation suggests that the worse the somatosensory precision of an external touch when participants moved their receiving hand, the worse the sensory precision for a self-generated touch during the same movement of the receiving hand. As the JNDs significantly increased only in these two conditions and these increases correlated with each other, this result provides evidence that their common experimental denominator, i.e., the movement of the left limb that receives the touches, was responsible for the increase in the JND.

A common finding in tactile gating studies is that the gating effects, both behavioral and electrophysiological, are stronger with higher movement velocities (*35, 38, 41, 44, 78*); the faster the limb movements, the worst the perception of the moving limb (see also (*79*) for findings in macaques). Therefore, one could hypothesize that any differences observed between the *gating* and *attenuation&gating* conditions might be due to differences in the velocity of the participants’ movements. Since no significant JND differences were observed between these two conditions (**Fig. 2**), which was further supported by Bayesian analysis, this concern can be excluded. However, one can argue that the PSE in the *attenuation&gating* condition was lower than the PSE in the *attenuation* condition because the participants moved faster in the *attenuation&gating* condition and not because of the reafferent nature of the touch. This concern can also be excluded since we observed that the participants moved slightly slower rather than faster in the *attenuation&gating* condition (20.3 *±* 0.003 cm/s) than in the *gating* condition (23.3 *±* 0.003 cm/s): (peak trial velocity; *t*(23) = -4.062, *p* < 0.001, *Cohen’s d* = -0.829, *CI*^*95*^ = [-0.004, -0.001], *BF*_*01*_ = 0.015). Although the total distances the participants ran with their left arm were comparable, participants in the *attenuation&gating* condition moved slower because they had to coordinate both their arms to tap the sensor that the left arm moves with the right hand (**Fig. 1d**). This difference was further confirmed when examining the peak velocities during the *test* tap between the two conditions (*gating*: 22.2 *±* 0.003 cm/s; *attenuation&gating*: 19 *±* 0.003 cm/s) (**Supplementary Fig. S5, S6 and S7**). Therefore, these kinematic differences do not explain the significant decrease in the PSEs in the *attenuation&gating* condition compared to the *gating* condition, but they actually underscore the importance of reafference in somatosensory attenuation.

## Discussion

The present study contrasted the conditions of attenuation and gating in a single experimental paradigm to investigate the relationship between the two phenomena on the same limb. Therefore, we independently manipulated the origin of the touch (reafference *vs*. exafference) and the state of the receiving limb (movement *vs*. rest), studying all four possible combinations of these levels. We replicated the classic phenomenon of predictive attenuation of touch (*1, 2, 17, 3*–*10*) by showing that somatosensory reafference feels weaker than somatosensory exafference. Importantly, however, this decrease in the perceived amplitude (PSE) was not accompanied by a concomitant worsening of somatosensory precision (JND). That is, participants had the same discrimination capacity (JND) for both reafferent and exafferent touches applied on their passive limb, a conclusion supported also by Bayesian statistics. Nevertheless, when the limb that received the touches moves, this voluntary movement *per se* led to a decrease in somatosensory precision (JND) for both reafferent and exafferent touches, replicating the classic tactile gating phenomenon (*35*–*39, 41, 43, 44*). The two effects did not correlate and did not interact but were summed when present together.

We must emphasize that in all experimental conditions, we measured the participants’ somatosensory perception on the same limb (i.e., the left limb). Although the participants moved their right hand to generate the tap on their left index finger in the *attenuation* and *attenuation&gating* conditions, our outcome measure was the perception of the (self-generated) tap on their left index finger. This approach is a particular strength of the current experimental design because it allows to independently induce, and separately study, the two phenomena on the same limb.

The main conclusion of the present study is that the predictive attenuation of touch and tactile gating are two distinct perceptual phenomena. Our findings can help conciliate several previous observations on gating and attenuation that have been studied in isolation in recent decades. First, attenuation is observed not only on the active limb (*8*) but also on a passive limb (the left hand in the present study), as long as the contact between the body parts is predicted by voluntary movement (*2*–*7, 9, 10, 14, 17*). In contrast, abundant evidence has shown that external touches applied to the limb contralateral to the limb that moves are not gated (*36, 47, 48, 50, 80, 81*). Second, a touch that results from a passive movement (*10*) or touches that are simultaneously presented in both hands (double touch) (*71*) are not attenuated. In contrast, gating effects have been repeatedly documented for passive movements, both electrophysiologically (*50*) and behaviorally (*36, 43, 76*). Third, self-generated tactile signals are attenuated as long as they are presented at the timing predicted by the action (*1, 2, 4*); even a 100 ms delay between the movement and its tactile feedback substantially reduces the attenuation of the latter. In contrast, externally generated stimuli are gated with less temporal sensitivity; for example, gating is observed for stimuli presented several (unpredicted) times during movement (*50*), at movement onset (*45*), and importantly, even hundreds of milliseconds before movement onset (*41, 43, 45, 47*). Fourth, although gating shows no specificity for the type of motor activity and manifests both during isotonic (*35*–*37, 48*) and isometric (*37, 81*) contractions (see also (*82*) for findings in macaques), attenuation is motor command-specific; a consistent but arbitrary and unnatural mapping between the motor command and the touch, for example, moving a joystick with one hand to produce touch on the other, does not produce attenuation (*5, 6, 73*). These results are not contradictory based on our findings; instead, they refer to different perceptual phenomena.

Based on this dissociation between gating and attenuation, we propose that findings from earlier studies using unimanual movements might reflect the combination of attenuation and gating effects because they have not isolated the two phenomena. For example, during a unimanual movement toward a target, Voudouris and Fiehler (*83*) showed smaller suppression effects at the time point of the maximal speed but increased suppression effects close to the time of contact (approximately 0-50 ms before contact) when the hand decelerated. Similarly, Fraser and Fiehler (*39*) showed greater suppression during the later phases of reaching, when the limb decelerated to approach the target (approximately at 140-180 ms before contact). In principle, these effects contradict the findings of Cybulska-Klosowicz et al., who showed that gating effects are greater during fast speeds than during slower speeds of elbow extension (a nonreaching movement) (*44*). According to Cybulska-Klosowicz et al., Voudouris and Fiehler (*83*) and Fraser and Fiehler (*39*) should have observed greater gating effects during peak velocity than during later stages. However, predictions of the somatosensory consequences of the movement (i.e., touching the target) during the later stage of reaching movements exist, and thus, one should observe somatosensory attenuation effects as well. Indeed, previous studies have shown significant attenuation effects at times very close to the time of contact (e.g., +/-150 ms around the button press) (*2, 4*). Based on our findings that attenuation and gating are distinct perceptual phenomena, we speculate that the findings of Voudouris and Fiehler (*83*) and Fraser and Fiehler (*39*) during the later stages of reaching movements reflect a combination of reduced gating effects because the speed of the movement decreases and increased attenuation effects due to the predicted consequences.

Concerning this point, active tactile exploration, such as when we move our hand to explore the shape or texture of an object (*84*), includes self-generated touch (contact between our finger and the object) experienced on the same limb that performs the voluntary movement. In contrast to the common view that active touch enhances performance during exploration, several studies have shown that texture perception with active touch is *not superior* to texture perception with passive touch (e.g., (*85*–*88*)). These findings led to the proposal that the relative motion between the hand and the object matters and not whether the subject actively moves the hand against the object/texture or the object/texture is moved against the passive hand (*88*). Having established that active touch is not superior to passive touch, the question that immediately arises is: Why is active touch not *inferior* to passive touch given the gating and attenuation effects present during voluntary movement? Importantly, studies have shown that when exploring textures in an unconstrained manner, we perform highly stereotyped exploratory movements (*89*) with slow mean scanning speeds (e.g., 52 to 120 mm/s (*90*)), which are not sufficiently fast to elicit significant gating effects (*44*). Similarly, we apply weaker forces when asked to explore the roughness/slipperiness of the texture than its hardness (*90*), and evidence has shown that attenuation is smaller for weaker forces than for stronger forces (*14*). We therefore propose that during tactile exploration (e.g., braille reading), we optimize the speed and pressure of our scanning movements to minimize tactile gating and attenuation effects and maximize the quality of the extracted information during tactile search (*44*).

Our study is the first to behaviorally dissociate the predictive attenuation of touch and tactile gating. A previous study (*91*) reported that the perceived intensity of an electrical stimulus applied to the tip of the relaxed left index finger does not change if this finger receives a self-produced force generated by the right index finger. However, in that study, the perception of touch was tested on a resting limb only, which is already known not to elicit gating effects (*36, 47, 48, 50, 80, 81*). Therefore, the two phenomena were not disentangled because they were not tested on the same limb, as in the current study. Similarly, another study (*67*) measured electrophysiological responses (somatosensory evoked potentials) to external electrical stimulation on the right and the left wrist while the participants produced a self-generated force with their right index finger on their left index finger. The authors did not measure the perception of external touches during movement (gating) but only the perception of self-generated touches during rest (attenuation) and thus were unable to dissociate the two phenomena behaviorally. Moreover, the authors observed modulation of the late electrophysiological responses depending on whether the participants produced a self-generated force, and they suggested that the attenuation and gating might share the same mechanism of reduced sensory precision. However, as shown in the present study, sensory precision is reduced during gating and not during attenuation, suggesting that a common mechanism governing both phenomena is unlikely.

Motor control relies on integrating afferent sensory information with efferent motor signals (*92*). Distinguishing between gating and attenuation is fundamentally important for motor control theories because it may indicate a different integration or weighting mechanism of the motor and sensory information, depending on the context. During voluntary motor control, these basic processes most likely coexist and cooperate, but our study suggests that they are distinct processes. Interestingly, when the two phenomena were copresent on the same limb (*attenuation&gating* condition), the participants’ performance showed no interaction effects but simple additive effects of the *attenuation* and *gating* condition. Although this condition involved a bimanual movement (i.e., participants moved both hands) and one could hypothesize the need for increased attentional demands or the presence of interhemispheric inhibition mechanisms with respect to the other conditions, the two phenomena did not interact but simply added together.

One well-established computational framework inspired by engineering approaches posits that the brain produces motor commands through an inverse model (*93*) or controller (*94*). A copy of the motor command, termed ‘efference copy’, is used by a forward model to predict the expected sensory feedback of the movement, which is then combined with the actual sensory input to estimate the state of the body (*61, 93*–*95*). With respect to the attenuation of sensory reafference, it has been proposed that the prediction signal of the forward model is used to ‘cancel’ the sensory reafference (*18, 61, 91, 96, 97*). In other words, central motor processes play a more important role in somatosensory attenuation than actual sensory feedback. Support for this comes from studies showing that conditions that present highly predictable touches but in the absence of movement do not yield attenuation (*2, 10*) (see also (*98*–*100*) for similar conclusions). The dependence of attenuation on action prediction was further shown when participants attenuated the touches applied to one hand that were predicted by their other hand’s movement, even when the two hands unexpectedly failed to make contact (*3*). Providing further confirmation, neuroimaging studies on somatosensory attenuation consistently report activation of the cerebellum (*12*–*16*), a structure that is associated with motor prediction (*92, 101*–*104*). Our findings of a reduced perceived magnitude of somatosensory reafference compared to exafference (i.e., lower PSEs in the *attenuation* and *attenuation&gating* conditions compared to the *baseline* and *gating* conditions, respectively) support this forward model mechanism for the attenuation of self-generated input.

In contrast, this computational account that relies on action prediction, efference copy, and internal forward models is not applicable to tactile gating since gated touches can be of an exafferent nature and occur at any (unpredictable) time during movement, even before movement onset (*41, 42, 52, 76*). In other words, no information is available that the brain can use to predict exafferent touches with the forward model because no causal relationship exists between the motor command and sensory input. This observation is consistent with the proposal that peripheral afferent signals from muscle spindles and joint afferents play the major role in gating (*41, 50*) and that gating effects have also been observed during passive movement, without significant differences in active movements (*36, 76*) (but see (*79*)). Then, if efference copy is not the basis for gating, how are the gating effects computationally explained?

The alternative computational framework of active inference has been proposed to explain somatosensory attenuation (*65*); however, this framework was based on an assumed equivalence between attenuation and gating. The active inference approach refutes the necessity of an efference copy and emphasizes the importance of a generative model and reflex arcs in the place of forward and inverse models and controllers (*105, 106*). According to the active inference account, the brain predicts the sensory input that would be expected from a specific action, and the body moves to fulfill these sensory predictions. Motor commands are thus conceptually replaced by proprioceptive predictions, and action occurs as a way to minimize the proprioceptive prediction errors when the movement has not yet been executed (*107*). A major role is assigned to the precision (i.e., reliability), which weights these sensory prediction errors depending on the context and can be manipulated through attention allocation. Within this computational architecture, attenuation of somatosensory input is viewed as a reduction in the precision of somatosensory evidence during movement to allow the expression of proprioceptive predictions that trigger the movement (*65*). In other words, the agent attends away from all somatosensory inputs to execute the movement. However, this proposal does not address the attenuation of sensory reafference (self-generated touch) with respect to exafference (externally generated touch) since the agent should theoretically attend away from *all* somatosensory inputs during the movement, regardless of their source. In contrast, we observed that self-generated touch on a moving limb (*attenuation&gating* condition) is perceived as weaker than externally generated touch on a moving limb (*gating* condition). Moreover, the active inference proposal refers to the limb that moves, and thus, the framework might not explain the effects observed on a passive limb. Indeed, it is puzzling why increasing the precision of the proprioceptive prediction errors on the hand that is to move (right hand) would reduce the precision of somatosensory evidence on the contralateral limb that is not meant to move (left hand) and where there are no proprioceptive predictions. In contrast, the active inference account may sufficiently explain the tactile gating effect, i.e., the reduction in the precision of somatosensory input on the moving limb during movement. Our findings of a reduced precision of somatosensory input on the moving limb (i.e., higher JNDs in the *gating* and *attenuation&gating* conditions compared to the *baseline* and *attenuation* conditions, respectively) are compatible with this active inference mechanism for the gating of sensory input during movement.

Another mechanism that has been proposed to explain tactile gating is a backward masking or postdictive mechanism (*69*) that is not necessarily dependent on motor signals. Accordingly, when moving a limb, the sensation from the muscles, joints, and skin of the moving limb *masks* the externally generated touches that are applied on this limb. These sensations could potentially affect the perception of earlier stimuli in a postdictive manner (*76*), which might explain the gating effects observed for external touches applied even before the movement onset (*43*). Our gating findings of a reduced precision of somatosensory input on the moving limb are compatible with this backward masking mechanism. Furthermore, in the context of a passive movement (i.e., in the absence of motor commands), the sensations from the muscles, joints, and skin of the passively moving limb might mask the externally generated touches on the same limb, and this mechanism might account for the absence of gating differences between active and passive movements (*36, 76*).

Regarding the neural mechanisms underlying the two phenomena, it has been shown that somatosensory attenuation results in reduced activity in the secondary somatosensory cortex (*12*–*14*) and the cerebellum (*12, 14, 15*) and increased functional connectivity between the two areas (*14, 16*). Accordingly, the cerebellum predicts the sensory consequences of the action based on the efference copy and cancels somatosensory activity (*12, 14*). In contrast, sensory gating studies in primates have shown suppression effects at very early stations along the somatosensory pathway, including the spinal cord (*60*), the cuneate nucleus (*79*), and the thalamus (*108*). In support of our conclusions, Chakrabarti and Schwarz (*109*) observed suppression effects on rats at the level of the brainstem (at the first synaptic level), where motor predictions are unlikely available, and proposed that sensory attenuation and sensory gating must be distinct phenomena (*109*).

In another study, Ishiyama et al. (*110*) compared the neural responses of rats when they groomed themselves (as a model for self-touch) with those when the experimenter touched (or tickled) the rats (externally generated touch). The authors observed substantially inhibited somatosensory responses during self-touch compared to externally generated touch and tickling. Interestingly, the inhibition of responses during self-touch was observed in the somatosensory cortex in a widespread and global manner. The inhibition was not restricted to the somatotopically organized zones that were the target of afferent inputs from the body parts stimulated by self-touch, and similar widespread inhibitory responses were also observed for externally generated touches applied at the same time as the grooming action. Based on these findings, the authors proposed a global inhibition suppression model that does not distinguish between self- and externally generated touches. Based on our findings, we propose that these “global inhibition” effects might actually reflect sensory gating effects and not the attenuation of self-generated touches, as animals actively move their bodies when grooming themselves. Consequently, suppression effects are expected for both self-generated and externally generated touches because the animal is motorically active. However, since the authors did not record motor responses in any of the experimental conditions, the extent to which the reported effects are due to self-touch or movement is not clear.

Thomas et al. recently suggested that somatosensory attenuation is due to simultaneous tactile stimulation to both index fingers (*111*) (“double touch”) and not action prediction. However, this claim is not supported by earlier experiments. First, previous studies have shown that double touch is *not sufficient* to produce somatosensory attenuation. For example, when both index fingers are simultaneously tapped (bimanual stimulation) in the absence of any movement (*2*) or in the presence of a passive movement (*11*), somatosensory attenuation does not occur. Similarly, when participants perform the force-matching task or the force discrimination task (as in the present study) and thus receive bimanual tactile stimulation, but a distance or a spatial mismatch is introduced between their hands/fingers, the attenuation is substantially reduced (*7, 14, 112*) or not present (*6, 113*). Second, several previous studies have shown that double touch is *not necessary* for somatosensory attenuation. For example, attenuation is also observed during unimanual movements (*8*), or when tactile stimulation is only provided to the passive left index finger during the force-matching task and the participants are only imagining the action of the right finger (using kinesthetic-motor imagery) (*9*). Critically, Bays et al. (*3*) used a very similar setup to Thomas et al. to explicitly examine whether the double touch produces attenuation and observed attenuation even when the right index finger moved to touch the left index finger but unexpectedly missed the contact (*3*). Collectively, the above observations strongly suggest that bimanual stimulation (double touch) cannot explain the attenuation phenomenon and that action prediction attenuates the predicted touch.

Our results also contradict Thomas and colleagues’ proposal that attenuation is due to nonpredictive generalized gating processes (*111*). Our study clearly shows that attenuation and gating are different phenomena; if sensory attenuation was the same as tactile gating, *all* stimuli applied to the moving limb would be attenuated. In contrast, we show that sensory reafference is robustly attenuated compared to sensory exafference, both in passive and moving limbs. Similar results were reported in another study (*91*), where participants attenuated only their reafferent touches and not exafferent touches presented on the same limb simultaneously. Therefore, our results do not support the equivalence of attenuation and gating phenomena proposed by Thomas and colleagues (*111*).

Finally, Thomas et al. (*111*) showed that participants perceived expected touches as stronger than unexpected touches on the fingers of their left hand when these touches were triggered by moving the right index finger on the air (no touch on the right index finger). This finding led them to propose that action leads to an *enhancement* of the predicted touch rather than attenuation. However, these results must be interpreted with caution for the reasons described below. First, one must include a baseline condition where the participant does not move (*baseline* condition in the present study) to argue that a perceived sensation is enhanced or attenuated. Consequently, lower values than those in the baseline are considered attenuation, and higher values are considered enhancement. Thomas et al. did not consistently include such a baseline condition in their experiments, and therefore, it cannot be concluded whether there is an enhancement or reduced attenuation in one condition than in another. Second, Thomas et al. provided participants with an arbitrary mapping between the movement of one hand and sensory feedback on the other hand (e.g., lifting the right index finger delivers a touch on the left middle finger). Arbitrary mappings between movements and touch are known not to elicit somatosensory attenuation (*6, 7, 14, 91, 114*). Thus, the finding that nonnaturalistic conditions do not allow the formation of predictions and the attenuation of the produced touch is not surprising. Third, the study of Bays et al. (*3*) that used a similar setup as that of Thomas et al. and a separate control condition that Thomas did not include, showed attenuation and no enhancement of the predicted touch. Finally, a key argument of Thomas and colleagues is that expectations should amplify our self-generated sensations to make our experiences more accurate in the presence of sensory noise. As we described above, several experiments have shown that active touch does not lead to better performance than passive touch and that we move our digits in a way to maximize the information we can extract when exploring the tactile world around us. It is important to acknowledge that the enhanced perception of self-generated touch suggested by Thomas and colleagues (*111*) would not be a more precise perception, as participants would be experiencing the stimulus as more intense than it truly is. This type of perceptual bias would be inaccurate, similar to the attenuated perception that we register in the current study.

We conclude that the human brain uses two different basic processes to suppress reafferent and exafferent information during movement and rest. This separation of attenuation and gating may explain why we tense our muscles when being tickled by others to decrease our sensitivity to external tickles, although we cannot tickle ourselves because we attenuate our self-tickles.

## Materials and Methods

### Participants

After providing written informed consent, twenty-four participants (12 women and 12 men, 22 right-handed, 1 ambidextrous, and 1 left-handed) aged 21-40 years participated. Handedness was assessed using the Edinburgh Handedness Inventory (*115*). The sample size was set to twenty-four (24) before data collection commenced based on our previous studies using the same methods (*4, 10*), while ensuring a counterbalanced order of conditions. Three participants were excluded because of technical issues with the kinematic recordings and replaced by three new participants to reach the target sample size. The Swedish Ethical Review Authority (https://etikprovningsmyndigheten.se/) approved the study (no. 2016/445-31/2, amendment 2019-04536). All participants provided written informed consent.

### General Procedure

Participants sat comfortably on a chair with their arms placed on a table. Their left hands rested palm up, with their index fingers placed on a molded support. The right arms rested palm down on top of a set of sponges. In each trial, a motor (Maxon EC Motor EC 90 flat; Switzerland) delivered two taps (the *test* tap and the *comparison* tap) on the pulp of their left index finger through a cylindrical probe (25 mm height) with a flat aluminum surface (20 mm diameter) attached to a lever on the motor. A force sensor (FSG15N1A, Honeywell Inc.; diameter, 5 mm; minimum resolution, 0.01 N; response time, 1 ms; measurement range, 0–15 N) within the probe recorded the forces applied on the left index finger. Following the presentation of the two taps, participants were required to verbally indicate which tap felt stronger: the first or the second. A second identical force sensor within an identical cylindrical probe was placed on top of, but not in contact with, the probe of the left index finger (**Fig. 1**).

A wooden surface was placed under the motor and the sensors. This surface was placed on top of two commercially available drawer runners (IKEA, https://www.ikea.com/us/en/p/besta-drawer-runner-soft-closing-40348715/). One side of the runners was attached to the table with Velcro, and the other side was attached to the bottom side of the surface. With this configuration, the surface, with the motor, the sensors and the participants’ hands, could be moved forward and backward.

In the *gating* and *attenuation&gating* conditions (**Fig. 1c, d**), participants were asked to extend their elbow upon an auditory ‘go’ cue. The extension of the elbow moved the platform forward on the table (**Fig. 1f**). A piece of green tape on the table (**Fig. 1i**) indicated the start position of the platform, while a piece of red tape indicated the end position. The participants were asked to move the platform from the start position to the end position (distance = 25 cm). During the movement of the left arm, the participants received the *test* tap on their left index finger. Before the condition started, we emphasized to the subjects that their task was to pay attention to the force that they would receive during the movement rather than covering exactly the distance between the lines. Moreover, the participants were trained to perform the movement in approximately 1000-1500 ms after the ‘go’ cue and then stop. In the *gating* condition, the *test* tap was applied 800 ms after the ‘go’ cue to ensure that it was delivered during the movement. Similarly, in the *attenuation&gating* condition, the participants triggered the *test* tap during the movement. The *comparison* tap was applied 800-1500 ms after the test tap to ensure that the participants had stopped moving. Once the participants responded, they returned the platform to the starting position.

In all conditions, the *comparison* tap was delivered on the left index finger with a random delay of 800-1500 ms from the *test* tap. We opted to present the *test* tap before the *comparison* tap (fixed order design), consistent with previous studies (*2, 4, 10, 11*), to maintain the delay between the two taps constant across conditions and remove any effect of the comparison tap on the tap participants had to perform with their right index finger. For example, if the participants first received a 3 N *comparison* tap, they might press stronger to generate the subsequent *test* tap. A fixed order design might introduce a temporal bias to the participants (e.g., participants perceive the second tap as stronger), but any of these biases cancel each other in the comparisons between conditions.

In the *attenuation* and *attenuation&gating* conditions (**Fig. 1b, d**), the tap of the participants’ right index fingers on the force sensor triggered the *test* tap on their left index finger with an intrinsic delay of ≈ 36 ms. In these two conditions, participants were asked to tap, neither too weakly nor too strongly, with their right index finger, “as if tapping the screen of their smartphone”. This instruction was provided to ensure that the relationship between the force they applied with their right index finger on the force sensor and the force they received on their left index finger by the motor (2 N) remained constant throughout the experiment, thereby establishing perceived causality (*91*).

A motion tracking sensor (6DOF Polhemus Fastrak, USA, weight = 9.1 g, dimensions = 2.29 cm x 2.82 cm x 1.52 cm) was placed on top of the platform to record the motion of the platform due to the movement of the participants’ left arm. The sensor recorded the x, y and z positions at a sampling rate of approximately 120 Hz.

Each condition included 70 trials. The test tap was set to 2 N, while the intensity of the comparison tap was systematically varied among seven different force levels (1, 1.5, 1.75, 2, 2.25, 2.5 or 3 N). Each tap lasted for 100 ms. In every trial, participants verbally indicated which tap on their left index finger felt stronger: the first (*test*) or the second (*comparison*). Participants were told not to try to balance their responses (50% first and 50% second), and they were further instructed to make their best guess if the intensity of the two taps felt similar.

In addition, participants were administered white noise through a pair of headphones to preclude any sounds created by the motor to serve as a cue for the task. The loudness of the white noise was adjusted such that participants could clearly hear the auditory cues of the trial. In all conditions, the view of the pulp of the left index finger was occluded. Participants were asked to fixate on a cross placed on a wall 2 m opposite them, but they were allowed to look at the force sensor to guide the movement of the right index finger when needed (**Fig. 1b, d**). No feedback was provided to the participants about their responses.

### Force discrimination analysis

In each condition, the participants’ responses were fitted with a generalized linear model using a *logit* link function (Equation 1):

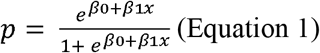

We extracted two parameters of interest: the PSE 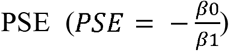, which represents the intensity at which the test tap felt as strong as the comparison tap (*p* = 0.5) and quantifies the perceived intensity, and the JND 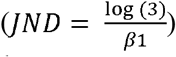, which reflects the participants’ discrimination capacity. Before fitting the responses, the values of the applied comparison taps were binned to the closest value with respect to their theoretical values (1, 1.5, 1.75, 2, 2.25, 2.5 or 3 N).

### Kinematic analysis

Both position and velocity data were smoothed with a moving average filter in MATLAB 2018a. Velocity was calculated as the first derivative of position. We calculated the minimum and the maximum position of the platform during the entire trial duration to calculate the distance participants moved in every trial under each condition. The peak trial velocity was defined as the peak velocity of the entire trial. The peak tap velocity was defined as the peak velocity during the period that the *test* tap was applied.

### Rejection of trials

After data collection, one hundred seventy-three (173) of 6720 trials (2.57%) were rejected. First, in thirty-four trials (34), the intensity of the *test* tap (2 N) was not applied accurately (*test* tap < 1.85 N or *test* tap > 2.15 N), and the responses were missing in sixteen (16) trials. Second, we rejected one hundred seventeen (117) trials from the *gating* and *attenuation&gating* conditions because participants either did not move their left arm (or moved it too slowly) during the *test* tap (mean velocity < 10 cm/s) or they moved it during the *comparison* tap (mean velocity > 5 cm/s). The thresholds were based on a previous study (*44*) showing no gating effects for velocities smaller than 5 cm/s. The analysis was therefore performed with 6547 trials.

### Statistical analysis

We used R (*116*) and JASP (*117*) to analyze our data. The data normality was assessed using the Shapiro–Wilk test. Depending on the data normality, we then performed planned comparisons using either a paired t-test or a Wilcoxon signed-rank test. We report 95% confidence intervals (*CI*^*95*^) for each statistical test. Effect sizes are reported as the partial eta-squared (η*p*^*2*^) values for the ANOVAs, Cohen’s *d* for t-tests or the matched rank biserial correlation *rrb* for the Wilcoxon signed-rank tests. In addition, a Bayesian factor analysis using default Cauchy priors with a scale of 0.707 was performed for all statistical tests to provide information about the level of support for the null hypothesis compared to the alternative hypothesis (*BF*_*01*_) based on the data. We interpret a factor between 1/3 and 3 as “anecdotal evidence” (*118*), indicating that support for either the preferred or null hypotheses is insufficient. Finally, correlations were determined by calculating the Pearson’s coefficient *r* because the data were normally distributed. All tests were two-tailed.

### Corrections for multiple comparisons

Since our PSE and JND comparisons were planned, we did not apply corrections for multiple comparisons. However, all results remained exactly the same when applying corrections for the false discovery rate (FDR) (*119*). In the correlation analyses, we corrected for multiple comparisons (FDR) since, although we expected correlations between the PSEs and between the JNDs, we had no *a priori* hypotheses for correlations between PSEs and JNDs.

## Acknowledgments

We thank M. Hauser, M. Chancel and L. Miller for their feedback on an earlier version of the manuscript. K.K. was supported by the Swedish Research Council (VR Starting Grant 2019-01909 granted to K.K.). H.H.E. and experimental costs were supported by the Swedish Research Council, Torsten Söderbergs Stiftelse, and Göran Gustafssons Stiftelse. We thank Kanaka Raghavan for assistance with the illustrations.

## Authors’ contributions

K.K. and H.H.E. conceived and designed the experiment. K.K. collected the data and conducted the statistical analysis. K.K. and H.H.E. wrote the manuscript.

## Supplementary Information for

**Fig. S1.**
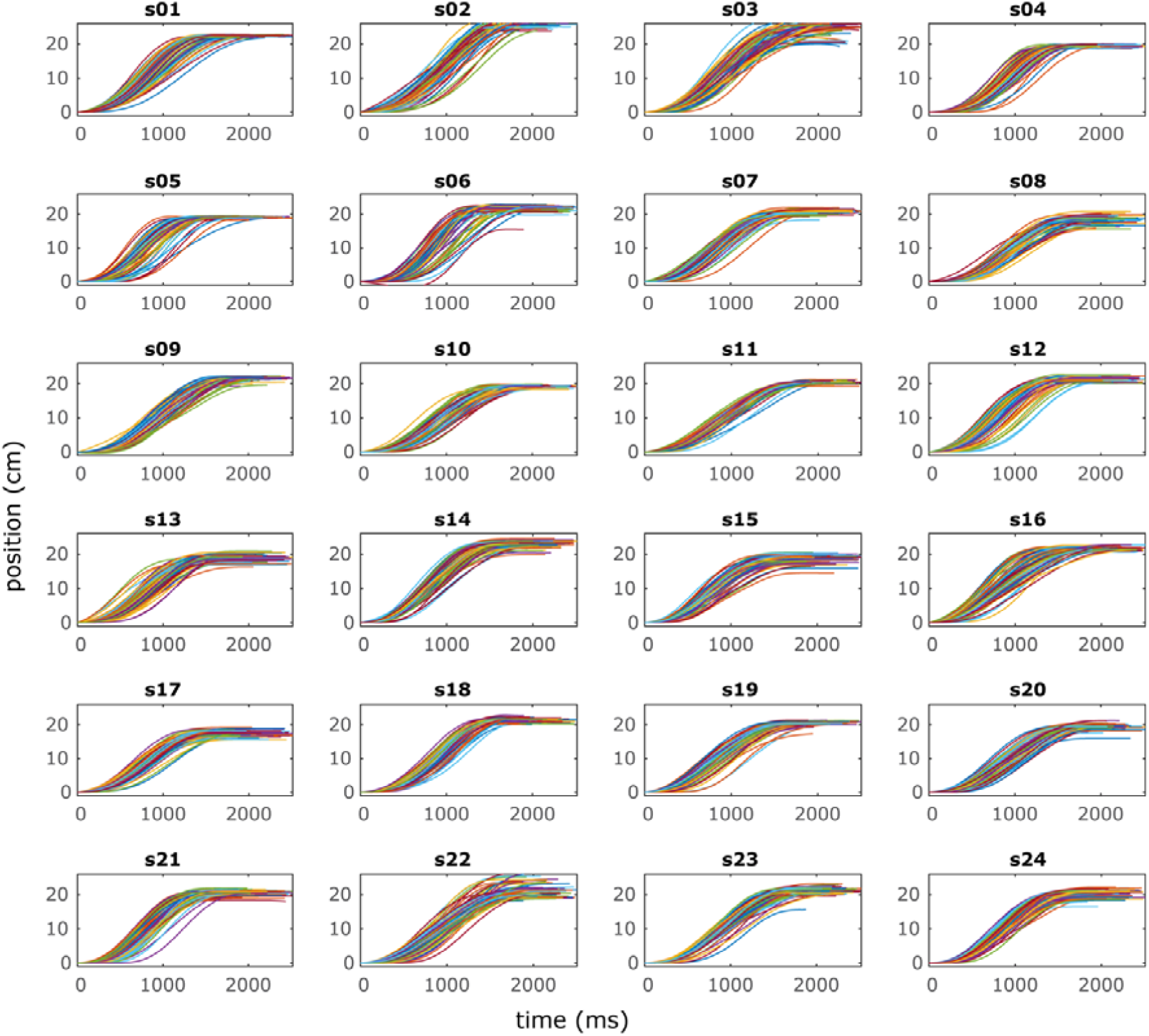
Position traces for the participants’ movements under the *gating* condition.

**Fig. S2.**
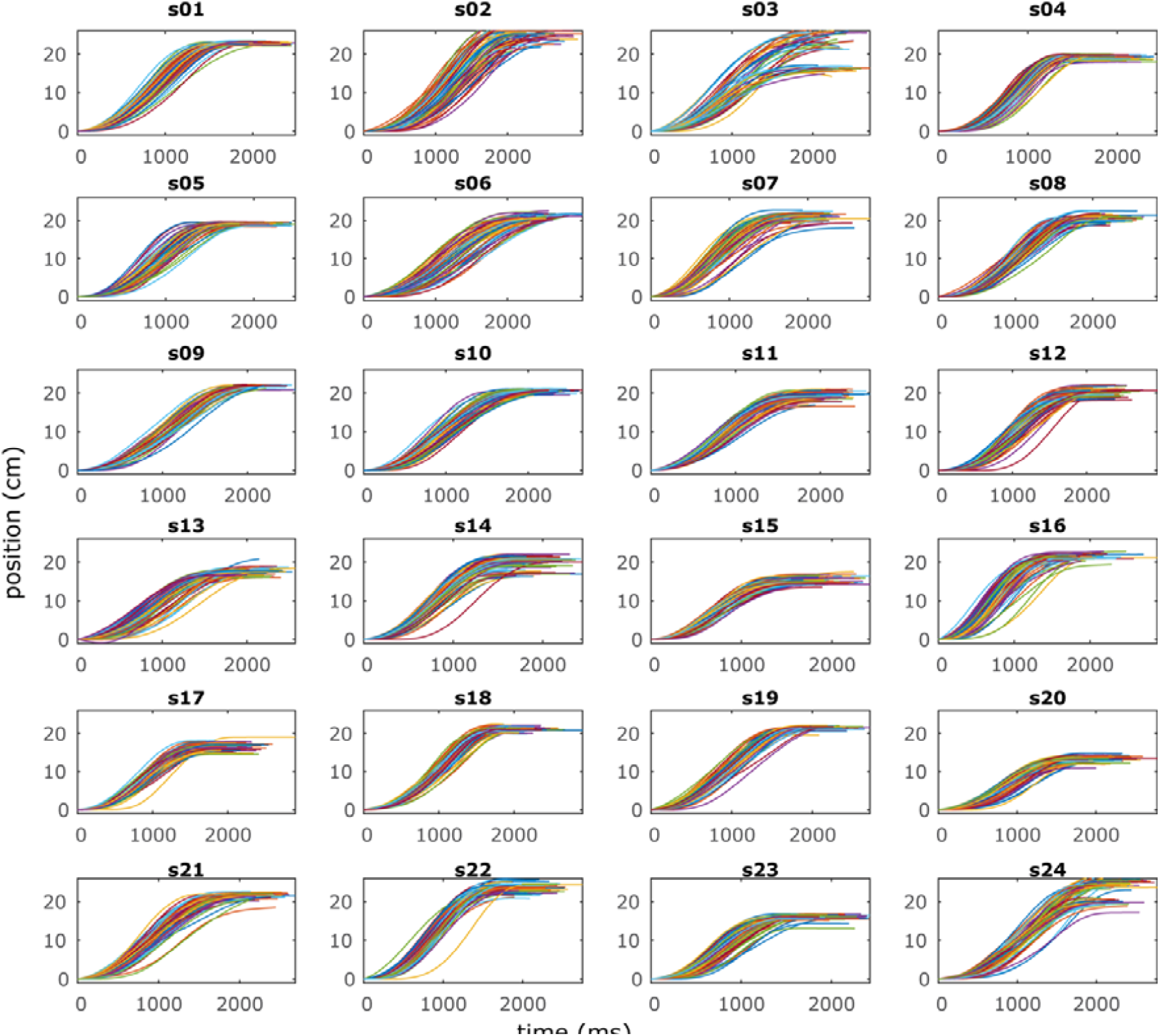
Position traces for the participants’ movements under the *attenuation&gating* condition.

**Fig. S3.**
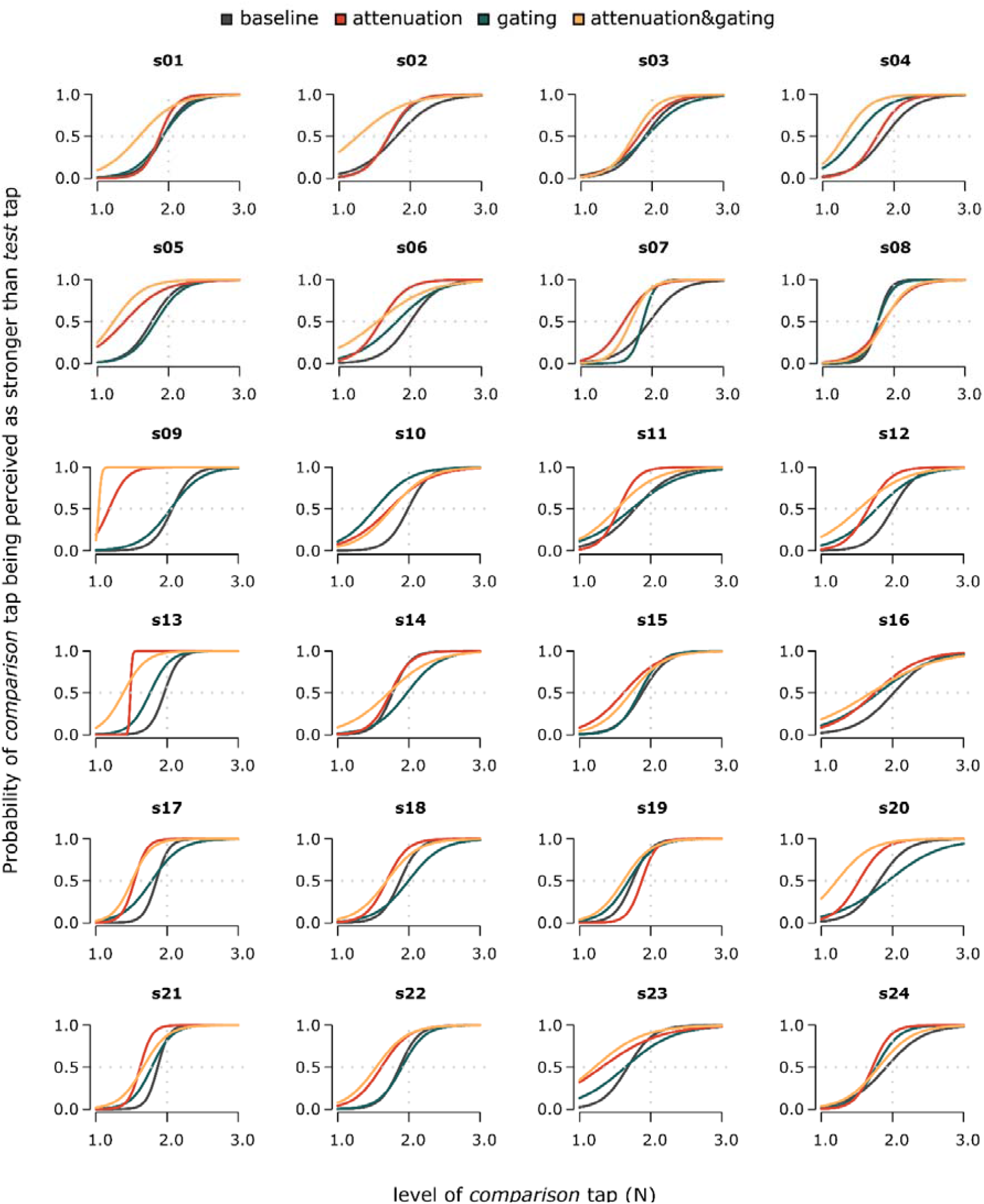
Fitted logistic models based on the participants’ responses under each condition.

**Fig. S4.**
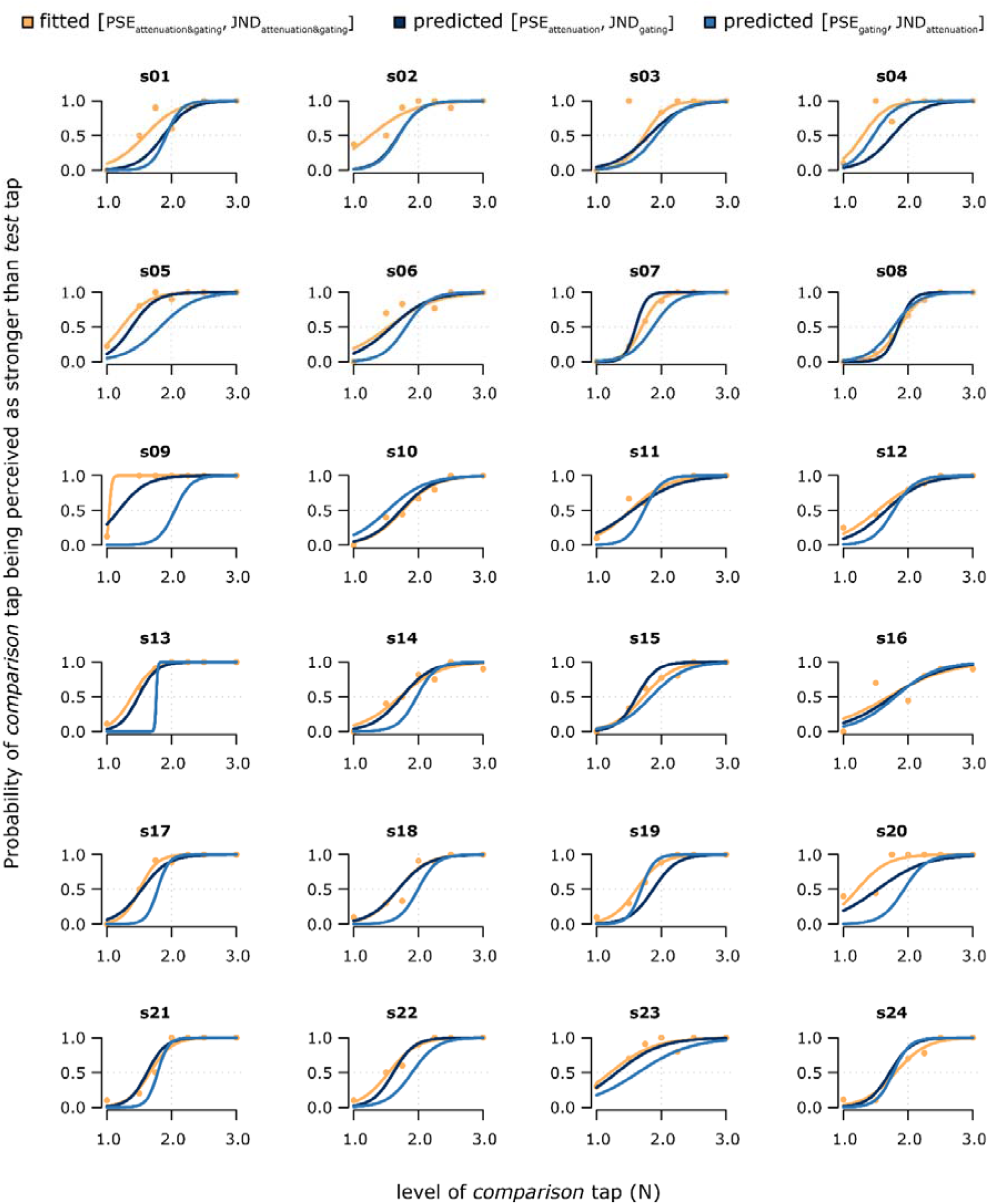
Fitted logistic models based on the participants’ responses under the *attenuation&gating* condition (yellow) and predicted logistic curves based on the participants’ PSE and JND in the *attenuation* and *gating* conditions (blue).

**Fig. S5.**
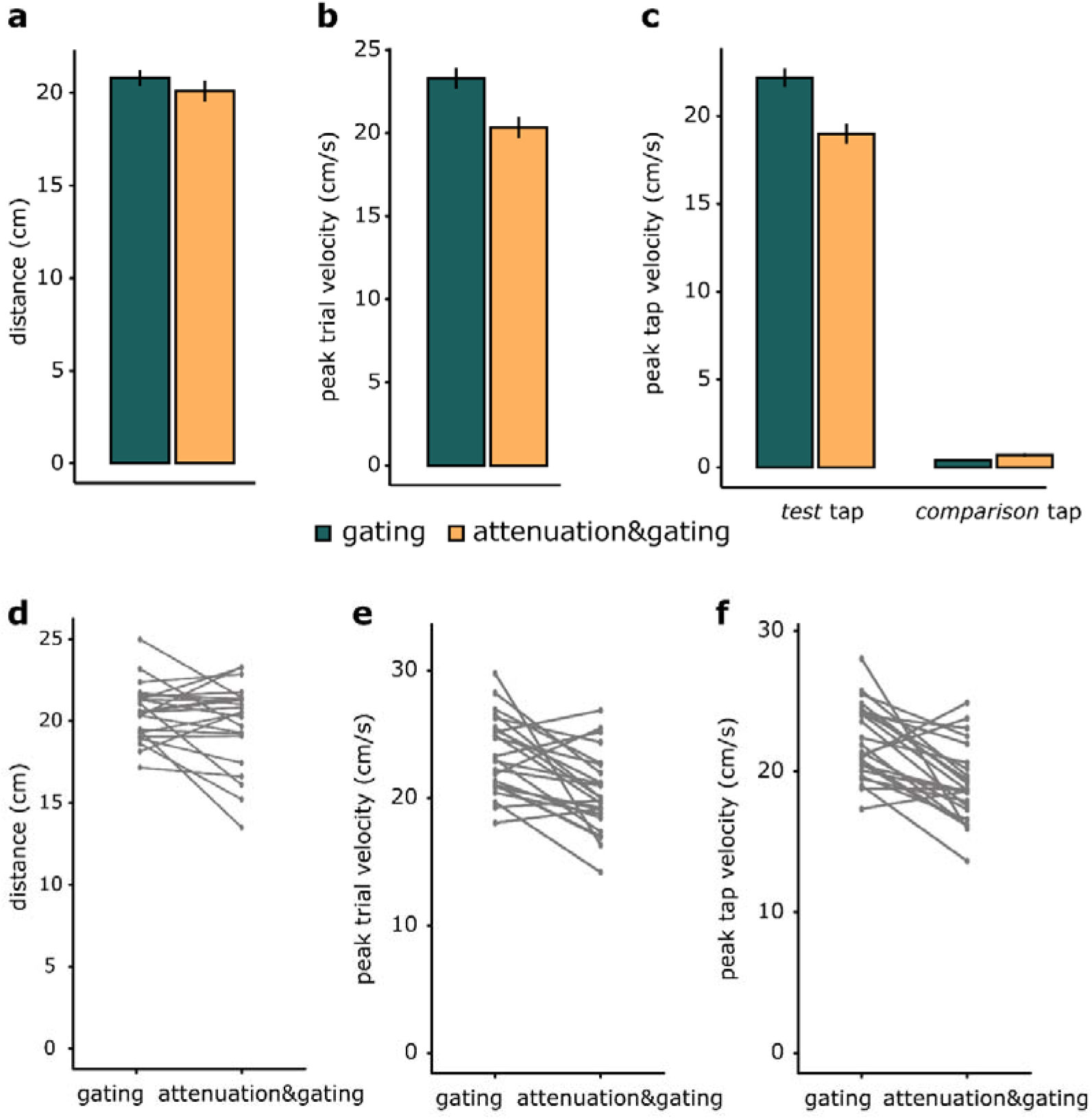
Movement parameters for the *attenuation* and *attenuation&gating* conditions. Bar graphs show **(a)** the total distance (mean ± SEM) run by the participants’ left arm, **(b)** the peak velocity (mean ± SEM) for the entire trial duration, and **(c)** the peak velocity (mean ± SEM) at the intervals of the two taps. (**d, e, f)** Line plots illustrate the differences in distance, peak trial velocity and peak tap velocity between the *gating* and *attenuation&gating* conditions.

**Fig. S6.**
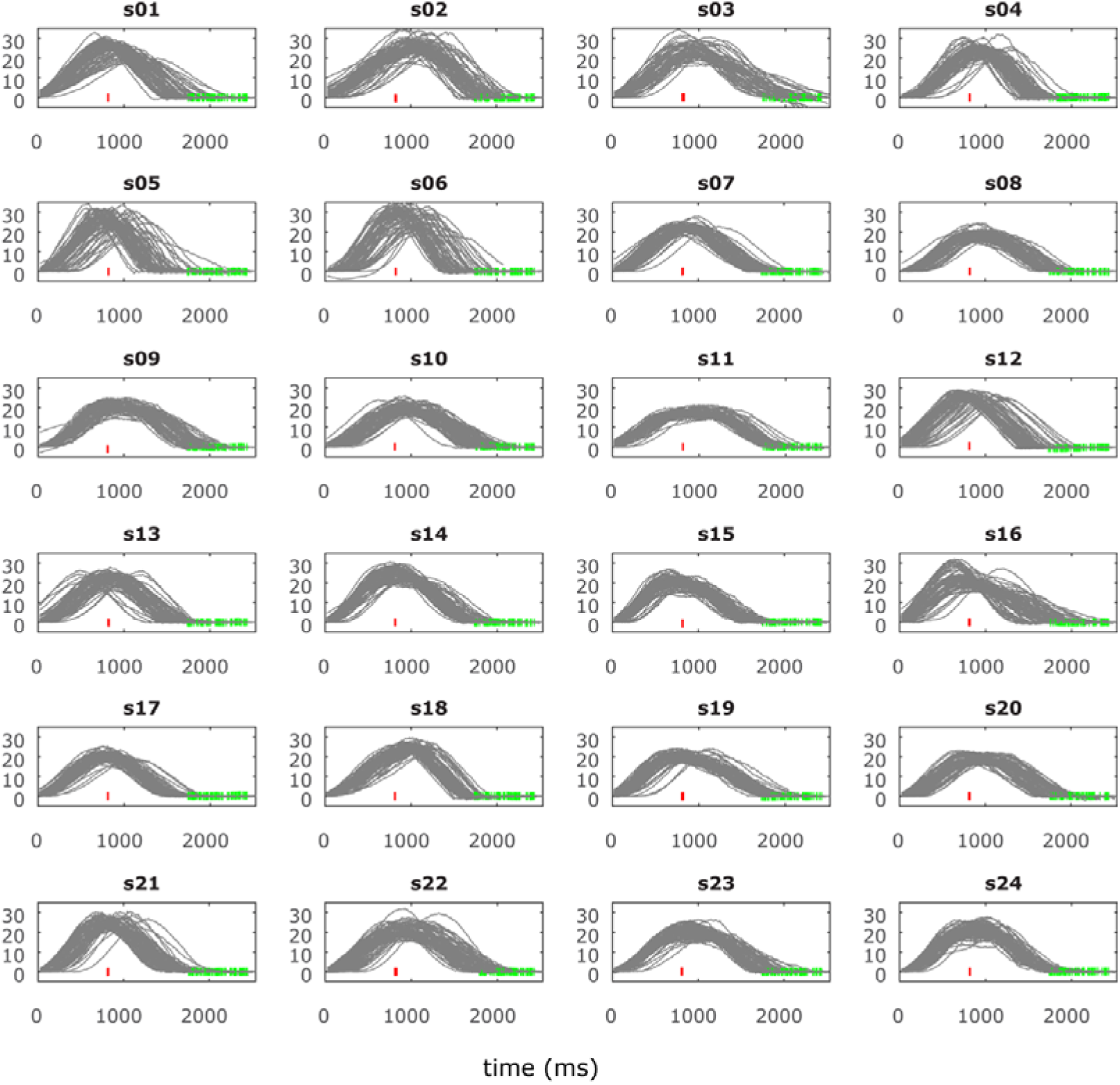
Velocity profiles for the participants’ movements under the *gating* condition. Red and green lines indicate the times when the *test* tap and the *comparison* tap were delivered, respectively.

**Fig. S7.**
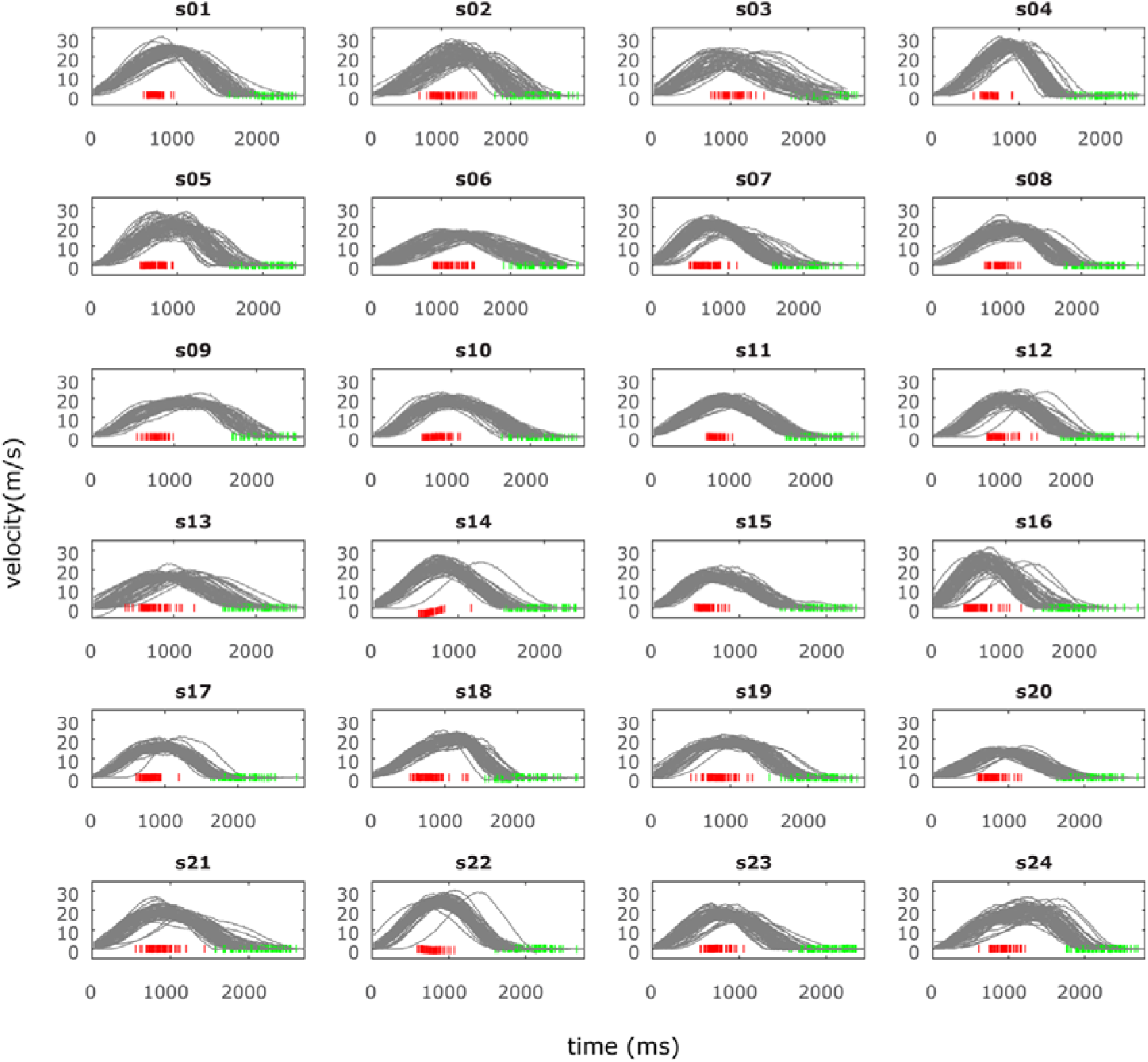
Velocity profiles for the participants’ movements under the *attenuation&gating* condition. Red and green lines indicate the times when the *test* tap and the *comparison* tap were delivered, respectively.

